# Modeling Molecular Mechanisms of Pirfenidone Interaction with Kinases

**DOI:** 10.1101/2024.03.22.586235

**Authors:** Prageeth R. Wijewardhane, Adrienne Wells, Matthew Muhoberac, Kai P. Leung, Gaurav Chopra

## Abstract

Scar formation is a process that occurs due to increased collagen deposition and uncontrolled inflammation. Previous studies have demonstrated that Pirfenidone (Pf), an FDA approved anti-inflammatory and anti-fibrotic drug can reduce inflammation *in vivo* as well as regulate activation of LPS-stimulated neutrophils. However, the molecular level mechanism of Pf’s action is not well understood. Here, we used neural networks to identify new targets and molecular modeling methods to investigate the Pf’s action pathways at the molecular level that are related to its ability to reduce both the inflammatory and remodeling phases of the wound healing process. Out of all the potential targets identified, both molecular docking and molecular dynamics results suggest that Pf has a noteworthy binding preference towards the active conformation of the p38 mitogen activated protein kinase-14 (MAPK14) and it is potentially a type I inhibitor-like molecule. In addition to p38 MAPK (MAPK14), additional potential targets of Pf include AKT1, MAP3K4, MAP2K3, MAP2K6, MSK2, MAP2K2, ERK1, ERK2, and PDK1. We conclude that several proteins/kinases, rather than a single target, are involved in Pf’s wound healing ability to regulate signaling, inflammation, and proliferation.

## Introduction

Pirfenidone (Pf) is an FDA approved anti-inflammatory and anti-fibrotic small molecule drug that is orally administered to treat mild to moderate idiopathic pulmonary fibrosis (IPF). Patients with IPF experience breathing difficulties as their lungs become scarred over time and ultimately become non-functional^1^. IPF occurs mainly due to repeated injuries followed by the inability of alveolar epithelial cells to effectively repair these injured wounds that leads to scar formation. Scar formation is a complex process that involves increased collagen deposition and prolonged inflammation. Whenever a tissue injury occurs, the innate immune cells (such as neutrophils and monocytes) infiltrate into the injured (wound) site. These innate immune cells influence wound healing through production of cytokines and growth factors that activate mitogen-activated protein kinases (MAPK) pathways, promote cell proliferation, protein synthesis, epithelialization, angiogenesis, as well as fibroblast proliferation and fibroblast-to-myofibroblast transition to regenerate tissue components such as collagen and other extracellular matrix proteins^2^. However, prolonged and uncontrolled inflammation may cause deregulated activation of wound healing cells, damaging the normal stages of wound healing that has been associated with excessive scarring such as scarring seen in IPF.

The fibrosis of lung and skin tissues share some common characteristics such as hardening, scarring, and overgrowing certain tissues and ascribes to the excessive deposition of extracellular matrix components like collagens. For an example, tissue samples taken from skin and lung lesions from Scleroderma or systematic sclerosis patients demonstrated a similar phenotype characterized by the presence of activated myofibroblasts, an indication of enhanced extracellular matrix synthesis and secretion of cytokines and chemokines^3–5^. Therefore, these similarities have enabled scientists to repurpose the anti-inflammatory and anti-fibrotic drugs such as Pf for treatment of scars seen after skin injury. However, the exact mechanism of action of Pf on treating IPF is not clearly understood. The estimated median time of survival from IPF is about 3 years^6^ and Pf has a half-life of 2.5 hours in healthy adults^6^. It’s reported that Pf can be extensively metabolized mainly by cytochrome P450 in humans^7^.The 5-methyl position in the molecular structure of Pf is the most vulnerable place to be metabolized and it can form 5-hydroxymethyl Pf and followed by 5-carboxylic acid metabolite^7^. Carboxylic acid metabolite is accounted for about 95 percent of the recovered Pf dose in adults^6^. Further, the rapid excretion of the Pf allows it to be used even three times per day for dose administrations in clinical trials and as a standard dosing schedule for IPF. According to previous studies of pulmonary and cardiac fibrosis, proliferation of fibroblasts and collagen synthesis can be inhibited by the Pf interference with growth factors that include the transforming growth factor (TGF-β) and basic fibroblast growth factor^8,9^. Moreover, Pf reduces the expression of inflammatory cytokines such as tumor necrosis factor (TNF-α), interleukin-4 (IL-4) and IL-13 by inhibiting the NOD-like receptor pyrin domain containing 3 (NLRP3) inflammasome. Further, the inhibitory effect mediated on the TGF-β is influenced by the inhibition of the MUC-CT phosphorylation by Pf^10^.

Our previous studies of Pf treatment *in vitro* and *in vivo* in the context of dermal application demonstrated that the drug reduces inflammation and fibrosis, suggesting its potential usefulness as a treatment for wound scarring. We found that Pf reduced the key pro-inflammatory mediators IL-1β, IL-2, IL-6, IL-13, G-CSF, and MIP-1α as well as decreased neutrophil infiltration in mouse deep partial-thickness (DPT) burn wounds^11,12^.Further, multiple studies suggested that Pf inhibited trans differentiation of human dermal fibroblasts (HDF) to myofibroblasts, weakened the contractile machinery of activated human dermal myofibroblasts and decreased collagen deposition and fibrosis-related gene expression^1,13^. Also, we found that Pf reduced p38-MAPK activation in TGF-β1-stimulated HDF and lessened chemotaxis, production of pro-inflammatory reactive oxygen species and cytokines (TNF-α, IL-1b, and IL-6) and degranulation of lipopolysaccharide (LPS)-activated human neutrophils^1,14^ as well as reduced activation of p38 MAPK, mitogen- and stress-activated kinase (MSK 1/2), protein kinase B (Akt), and c-Jun N-terminal kinase (JNK); and decreased pro-inflammatory mediators in porcine DPT burn wounds (unpublished data). However, per our knowledge, there is no study examining the molecular interactions of Pf with its target molecules that are essential in wound healing sequences and scarring of skin injury. Therefore, this study has extensively investigated the Pf molecule’s plausible action pathways using molecular modeling and machine learning methods (**Figure 1**) to understand how it regulates phenomena such as inflammation which leads to the attenuation of both the inflammatory and remodeling phases of wound healing for more optimal outcomes.

**Figure 1.**
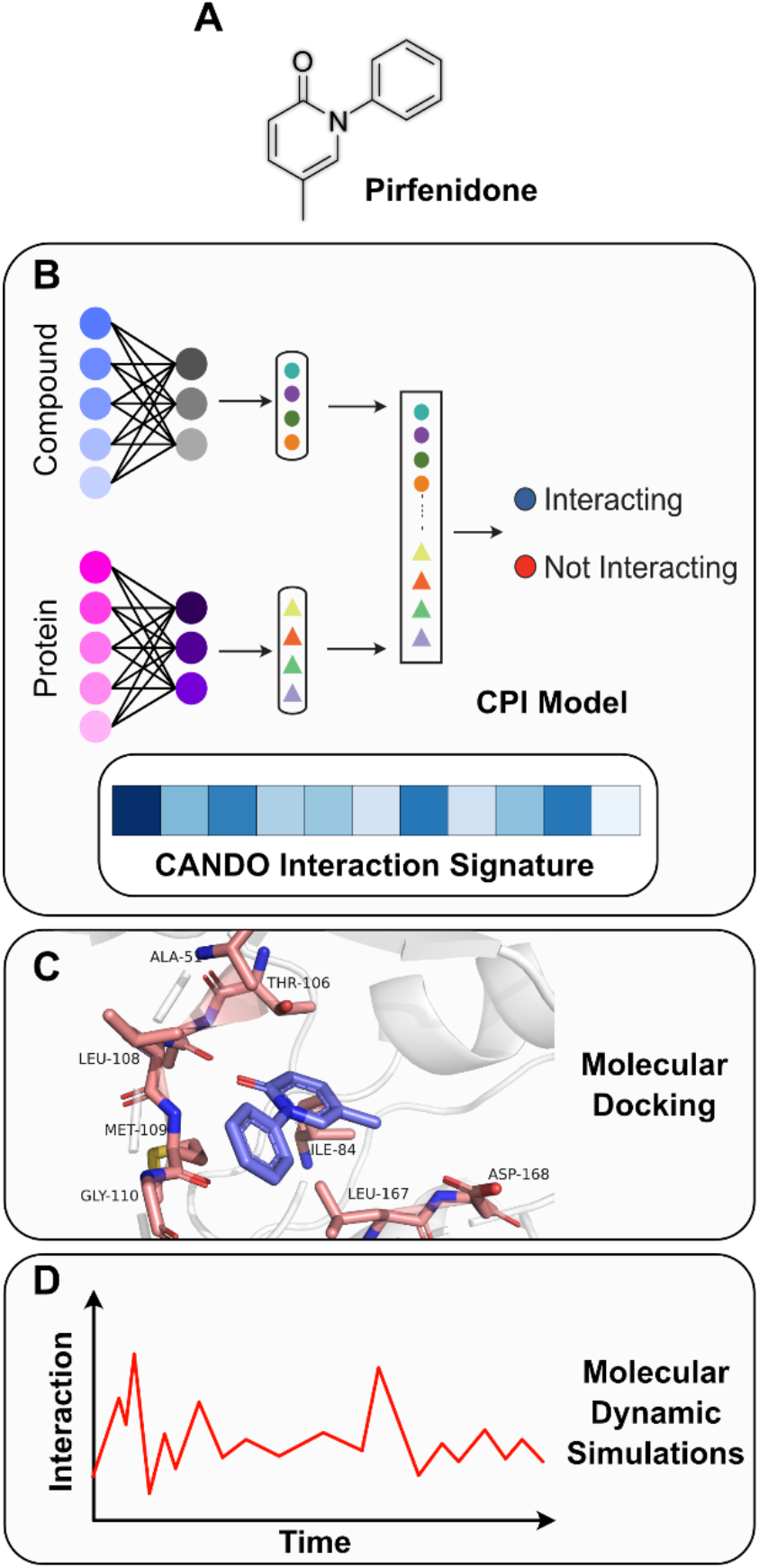
Overview of the machine learning and molecular modeling pipeline for Pf. (A) Chemical structure of Pf. (B) Compound-protein interaction (CPI) model and the CANDO signature to identify potential Pf targets. (C) Molecular modeling to analyze Pf interactions with selected targets. (D) Molecular dynamics simulation to investigate the time dependent stability of Pf-target interactions.

## Methods

### Neural Networks to Predict Pf Interaction with the Human Kinome and Other Proteins

Compound-protein interaction (CPI) machine learning model^15^ was used to identify potential kinase targets of the Pf small molecule. The CPI model utilizes two neural networks to predict the probability that an interaction between a specific compound and protein will occur^15^. We have also extended this approach previously by incorporating structural information to develop an energy-based graph-neural network^16^. In this work, the protein is represented as an amino acid sequence without any three-dimensional structural information provided to the model. The compound is initially input as a SMILES string and then is converted to a molecular graph prior to training the model. A convolution neural network is used on overlapping sequence motifs and a graph neural network is used on molecular graphs. Attention is used to determine interactions between individual sequence motifs and compounds. The weights of the individual neural networks are concatenated into a linear classifier which is used to predict the final binding probability. The CPI model used in this project was trained on the dataset with a near 50:50 split in positive and negative interaction (binding) data to identify binders from non-binders^17^. The model was trained to predict interactions between ATP and Pf with 536 human kinases and other proteins. The radius of the molecular fingerprinting algorithm used in the GNN was set to 2, the protein motif sequence length was set to 3 and each model was trained for 100 epochs.

### CANDO Platform to Predict Pf Interactions with the Human Proteome

We used, Computational Analysis of Novel Drug Opportunities (CANDO)^18^, implemented as a python package for analysis of drug-proteome and drug-disease relationships. CANDO can use thousands of proteins collected from public databases such as Protein Data Bank (PDB) to generate the interaction matrix with approved drugs and small molecules, most notably from DrugBank. An interaction scoring protocol has been used for the fast calculations of drug-protein interactions.

### Pf Binding Interaction Analysis with Molecular Docking

The CANDOCK^19^ docking software was used to perform all the docking analyses performed in this study. Binding sites of protein crystal structures were identified using the “find_centroids” function in the program. The cleaned 3D structure of Pf was prepared as a Tripos Mol2 file and the “prep_fragments” function in CANDOCK was used to determine which bonds in the molecule to be cut. Then, “make_fragments” function was used to produce PDB files for all the fragments produced in the previous step. Next, the software used the “dock_fragments” function to dock each and every prepared fragment at the predicted binding site of the protein of interest. Finally, the function “link_fragments” was used to link all the docked molecular fragments to form the original ligand structure with their respective docking score energy scores based on default parameters.

### Molecular Dynamics of Active and Inactive p38-MAPK with Pf and ATP

The time dependent interaction behaviors of Pf and ATP were analyzed by molecular dynamics (MD) simulations. The main reason to run ATP simulations is to use it as a control and compare the analysis with the same Pf simulation. We calculated the association time to quantify how much time as a percentage of the total simulation time the ligand is in close proximity of MAPK14 hinge region in the active site. These MD simulations were done using the GROMACS^20^ software package. The CHARMM36 forcefield^21^ was used for all simulations. A dodecahedron box shape was used to simulate the complex with TIP3P water solvation and neutralized total charge of the complex. Energy minimized complex was used to equilibrate with NVT, NPT ensembles, respectively. Then MD simulations were carried out for 1 µs time for the analysis based on the stability of ATP MD simulations as controls. Here, the active and inactive conformation classification was used based on protein phosphorylation. We used hinge region residues as a reference to calculate distance fluctuations between ligands and the protein to assess association times.

## Results and Discussion

### Protein structure selection for docking

Based on our previous studies^1,14^ we have identified available active and inactive structures of p38 mitogen-activated protein kinase targets (p38-MAPK), mitogen- and stress-activated kinases (MSK1/2), serine-threonine protein kinase (AKT1/2), and focal adhesion kinase (FAK) as potential Pf interacting candidates. Selecting both active and inactive structures were done to determine how the interaction affects the kinase signaling and to analyze whether it affects ATP binding of these kinases. First, kinase structures were preprocessed using our in-house molecular docking software, CANDOCK^19^, to determine “druggable” binding sites that includes the ATP binding site. P38-MAPK has four classes such as, p38-α (MAPK14), p38-β (MAPK11), p38-γ (MAPK12 / ERK6), p38-δ (MAPK13 / SAPK4). The selection was made for both human (*Homo sapiens*) and mouse (*Mus musculus*) proteins, for a total of 30 structures (**Table S1**) used for modeling, so that we can establish translation potential and changes during molecular modeling between the two species.

### Machine Learning and CANDO Analysis to Identify Additional Potential Targets of Pf

The distribution of binding prediction results using the compound-protein interaction (CPI)^15^ prediction model for both ATP and Pf against human kinases is shown in **Figure 2A**. Out of these predictions, several interesting targets came up as binders for Pf such as MAP3K4, MAP2K3, MAP2K6, MSK2, MAP2K2, ERK1, ERK2, PDK1 and details are tabulated in the **Table S2**. The CANDO^18^ interaction signature of Pf was calculated using 14,595 human proteins and the results are summarized as a histogram in **Figure 2B**. Interestingly, several targets appeared in this CANDO interaction signature of Pf such as cAMP-dependent protein kinase catalytic subunit alpha protein was also predicted by the machine learning approach. Using CANDO, the top three predicted protein targets for Pf are: macrophage migration inhibitory factor, caspase-3, and leukotriene A-4 hydrolase.

**Figure 2.**
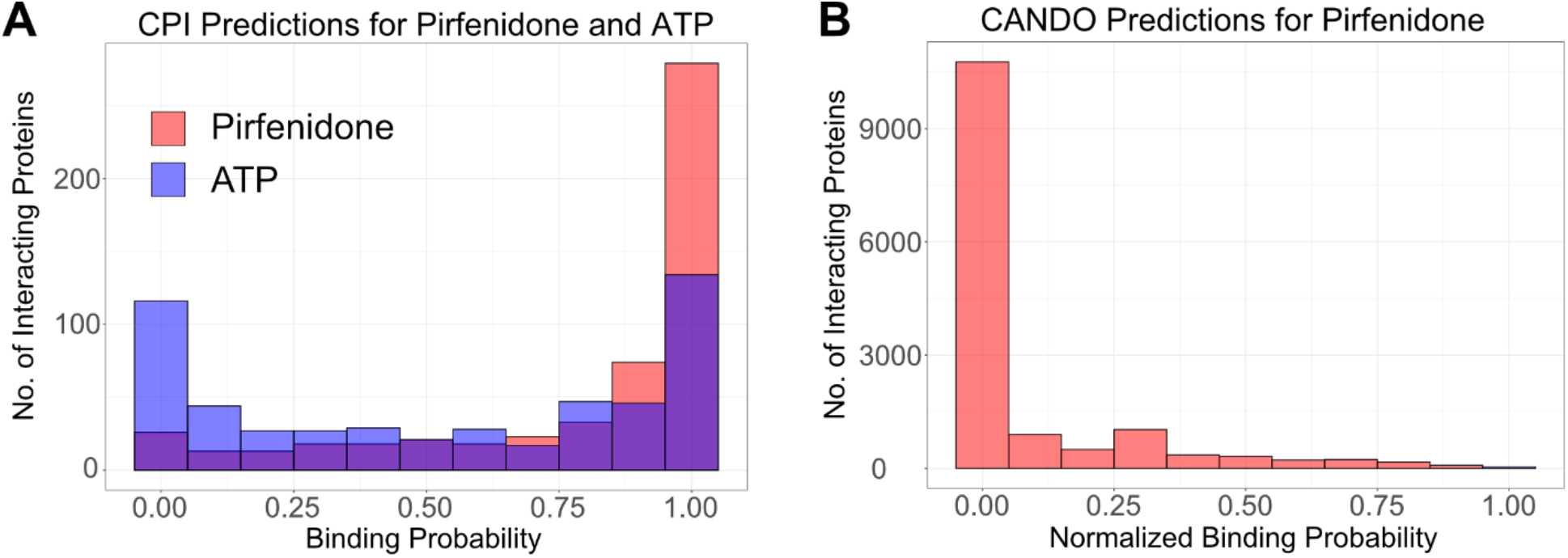
Compound-protein interaction (CPI) model and CANDO results for predictions of Pf binding. (A) Interacting proteins distribution of binding probability of ATP and Pf with human kinome and other targets (536 total) predicted using the CPI based graph-neural network model. (B) Normalized binding probability distribution predicted for Pf using the CANDO program.

### Molecular Docking to Analyze Pf Binding with Different Kinases

#### Selection of a Reliable Molecular Docking Score Function

To quantify the interaction of Pf in the presence of ATP, we first developed an interaction energy score with proper controls so that we can computationally test and quantify the effect of Pf in ATP binding sites. We used ATP as a “control” in our computational experiments to determine ATP binding based on inactive (ATP binders) and active kinases (ATP non-binders). This was done by docking ATP to all known crystal structures of active and inactive forms of p38-MAPK (**Table S1**) to determine the best energetic scoring function. CANDOCK^19^, our in-house docking software, was used for docking. Briefly, CANDOCK is a flexible docking method that breaks apart all bonds in the small molecule and assembles it inside the binding pocket using graph-based techniques, thereby using induced-fit as a mechanism to identify several conformations of small molecule fragments at different locations in the binding pocket. CANDOCK contains 96 different scoring functions to calculate the binding energy score. Using positive and negative training data based on ATP control binding with inactive and active kinase structures, we calculated Cohen’s kappa metric to identify the best scoring function. Cohen’s kappa provides an estimate of inter-model reliability to identify the scoring function and its reliability is better than random predictions for binding^22^. Since we were interested in estimating effect of Pf with p38-MAPK and other kinases, this calculation is essential to quantify the effect with ATP binders *v*ersus non-binders (training data). Here, we consider an active kinase form, when a protein is catalytically active or phosphorylated with converting ATP to ADP. We identified FMR15 scoring function^19^ in CANDOCK as the best scoring function with a kappa value of 0.41, indicating a moderate model for estimating binding of Pf in comparison to ATP.

#### Pf Binding with Active and Inactive Kinases

All the relative docking scores were calculated with proper controls for active and inactive targets. For inactive kinases, the Pf docking score is taken relative to the ATP docking score as ATP is the most preferrable towards the inactive conformation. However, in the active conformations of kinases, the Pf docking score is taken relative to the ADP docking score as it is the most preferrable form towards the active conformation. This is to make sure that the relative docking score is compared to the natural ligand of the conformation of the kinase. The analysis was carried out using the mean docking score of the top five poses of each docking result and summarized in **Figure 3A**. Relative docking score of Pf with active *vs*. inactive conformations of MAPK14 show that Pf has a preference towards the active conformation. This result agrees with the long molecular dynamics simulations which were done with MAPK14 structures as well. For another target, FAK, Pf did not show any preference out of active *vs*. inactive conformations. However, it showed a preference towards the active Akt1 conformation compared to inactive conformation based on the docking analysis.

**Figure 3.**
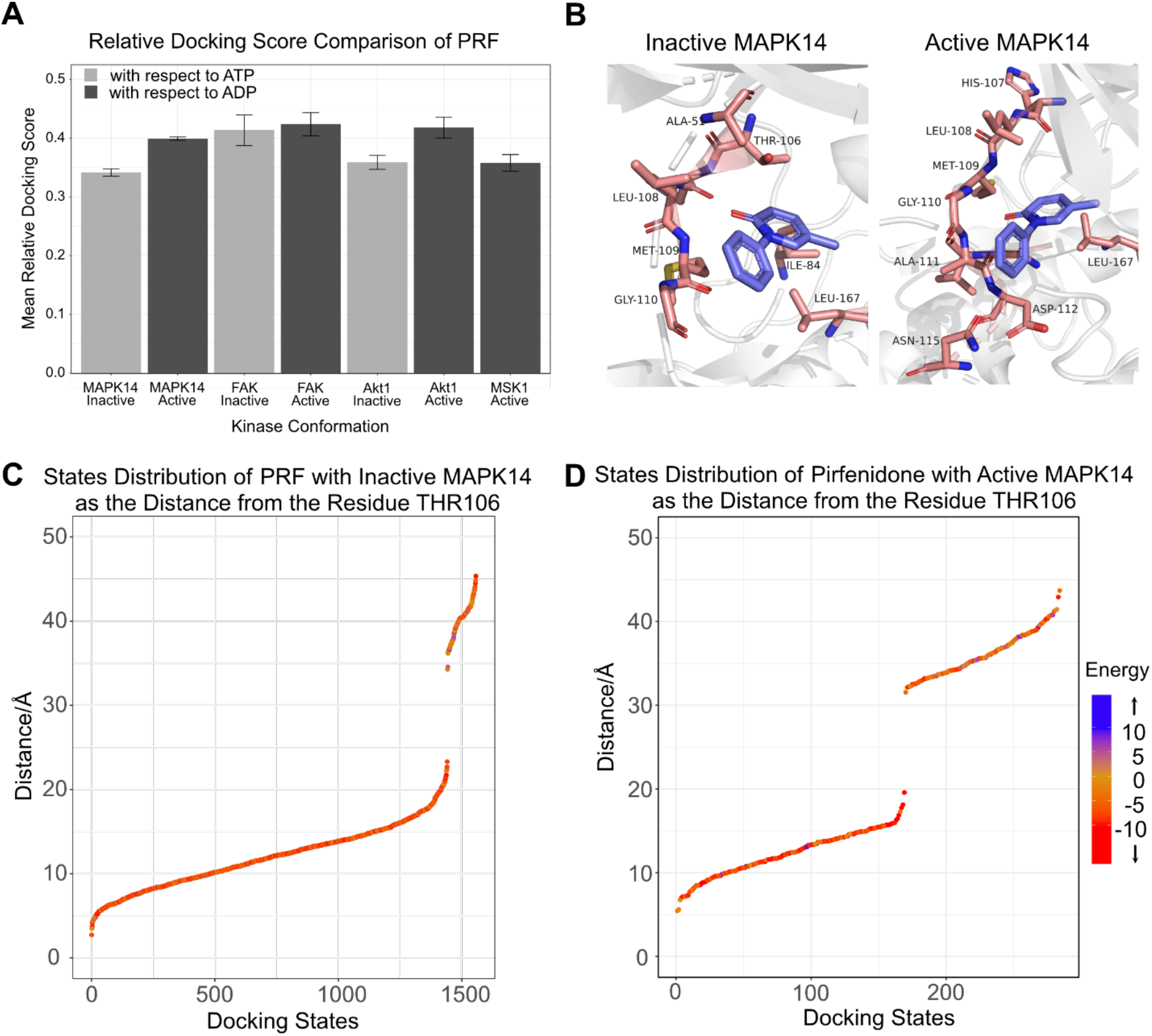
Molecular docking analysis of Pf with selected targets to understand its interactions at the binding site. (A) Mean relative docking score of with respect to the control. ATP is considered as the control ligand for inactive kinase conformations and ADP is used for active conformations. (B) Pf binding poses at inactive and active MAPK14 binding sites. (C) & (D) Docking state distribution along with the docking score energy of Pf with inactive and active conformations of MAPK14.

Moreover, starting with our docked structures, we used Discovery Studio Visualizer^23^ software with default parameters to analyze key interactions of ATP and Pf with MAPK14. Our analysis showed that most of the amino acid residues in inactive MAPK14 such as Thr-106, His-107, Leu-108, and Met-109 have formed similar interactions with both Pf and ATP. **Figure S1-A** shows these 2D interaction maps for a representative inactive MAPK14 crystal structure (PDB ID: 5ETI). Conversely, Pf and ADP interaction maps with the active conformation of MAPK14 (PDB ID:3PY3) are shown in **Figure S1-B**. Again, interaction maps indicate that amino acid residues of active MAPK14 such as Leu-108, Met-109, and Gly-110 form plausible common interactions with both Pf and ADP. Results suggest that even though the binding site is quite large, Pf prefers the area where ATP binds in both MAPK14 conformations.

It is well known that type I inhibitors bind with active kinases (DFG-Asp-in, αC-helix in) in the region occupied by the adenine ring of ATP^24,25^. Conversely, type-I½inhibitors are defined as inhibitors that bind to a DFG-Asp-in inactive kinase conformations and type-II inhibitors interact with DFG-Asp-out inactive kinase conformations also share the same area^26^. The residues 106 – 110 in MAPK14 are known as hinge region residues and it is reported that all type-I, type-I½and type II inhibitors interact with these hinge region residues^25,26^. The interaction maps of Pf with both inactive and active MAPK14 (**Figure S1**) show that it interacts with most of these hinge region amino acid residues. Given the similarity of Pf binding in these docking results, we cannot conclude whether Pf act as either type-I, type-I½or type-II inhibitor with MAPK14. Therefore, further investigation is needed to confirm the most probable binding mechanism. This is potentially exciting as it indicates that Pf may be functionally affecting signaling due to specific conformational change and performing molecular dynamics simulations could provide valuable insights for understanding this binding process.

Next, we investigated the areas of Pf binding on active and inactive conformations of MAPK14 proteins with molecular docking to investigate preferred bind sites. Pf binding poses of inactive and active MAPK14 crystal structures are shown in **Figure 3B**. Even though docked Pf resides around similar areas in both conformations, due to the conformational change in the active MAPK14, a beta sheet of the protein covers the Pf binding site from the top which allows the small molecule to buried inside of the binding site, potentially stabilizing its binding ability. **Figure 3C** and **Figure 3D** show distance plots for Pf docking from the first hinge region residue Thr-106, with active and inactive MAPK14, respectively. Each dot in the plots represents a single pose, and the color of the dot indicates the pose’s docking score energy. If the docking score energy is positive (blue color), it represents an unfavorable interaction between the small molecule and the protein. Conversely, if the docking score energy is negative and red in color, it indicates a favorable small molecule-protein interaction. Two distinctly separated scatter clusters in **Figure 3D** suggests that most likely there are two main Pf binding areas in the active MAPK14 conformation.

#### Molecular Dynamic Simulations to Understand Time-dependent Behaviors of Pf

Based on docking studies of both Pf and ATP, we selected the 19^th^ pose of Pf and the 20^th^ pose of the ATP for the MD simulation with inactive MAPK14. On the other hand, the top docked pose for ATP and 2^nd^ pose for Pf were chosen for the MD simulation analysis with the active MAPK14 conformation. These selections were made based on the presence of key interactions with the hinge region residues using both molecular docking and short MD simulations analyses. All the MD simulations were performed using the GROMACS^20^ software package. **Figure S2** shows that the fluctuation of distances of ATP from hinge region residues with time. Results of the completed (1 µs) simulation of ATP with inactive MAPK14 suggested that our control worked well since ATP showed short distance fluctuations from the hinge region residues. The analysis showed that the ATP stays within the 10 Å proximity of the hinge region residues Thr 106 (1000 ns, 100%), Leu 108 (924.95 ns, 92.5%) and Met 109 (1000 ns, 100%) with an average of 974.99 ns (97.5%) association time out of 1 µs.

Next, we studied the time dependent behavior of Pf with active and inactive MAPK14 proteins. **Figure 4A** shows the distance fluctuations with time of Pf from the hinge region residues in the catalytic binding sites. Based on the long 1 µs MD simulation for active MAPK14 compared to that of inactive MAPK14, we found that the Pf association results were the opposite between active *vs*. inactive conformation. Pf showed a better stability with the active conformation and higher association times out of the simulation time of 1 µs. The analysis showed association times of 1000 ns (100%) with Met 109, 1000 ns (100%) with Leu 108, and 999.53 ns (99.95%) with Thr 106 out of the 1 µs MD simulation. The average association time of Pf with active MAPK14 was 999.85 ns (99.98%) out of 1 µs.

**Figure 4.**
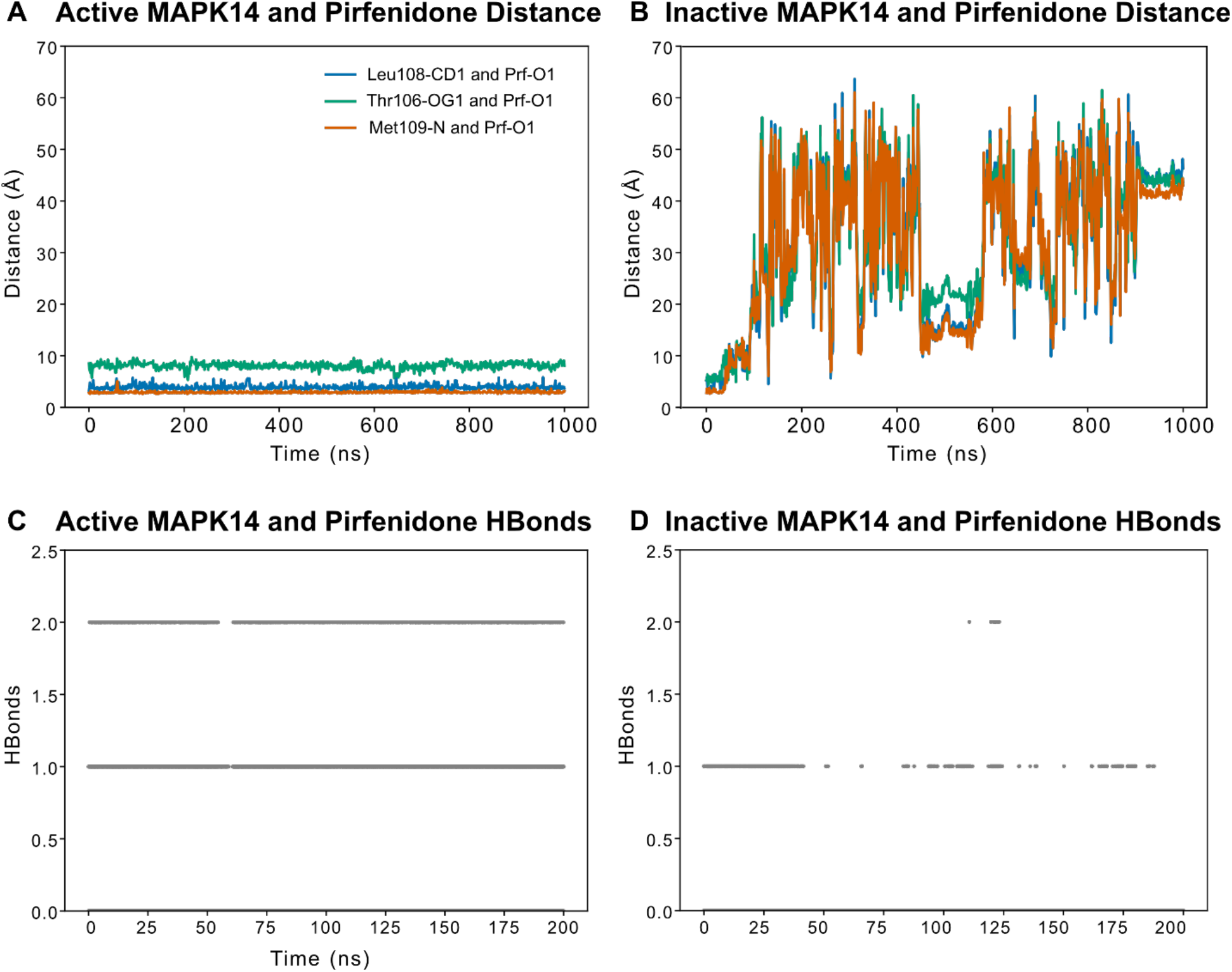
Molecular dynamics simulations results of Pf with active and inactive MAPK14 conformations. (A) & (B) Distance trajectories between different atoms on Pf and different hinge region residues on the MAPK14 for active and inactive MAPK14 conformations, respectively. (C) & (D) No. of hydrogen bonds formed between Pf and the two conformations of MAPK14 through out the first 200 ns of the simulation.

In contrast, a similar analysis showed that Pf did not stay within 10 Å radius around hinge region residues of inactive MAPK14. Association times observed were 75.28 ns (7.53%) with Met 109, 74.45 ns (7.45%) with Leu 108, and 70.07 ns (7.01%) with Thr 106. On an average, the association time was about 73.27 ns (7.33%) out of 1 µs (**Figure 4B**) suggesting that Pf does not prefer to bind to the inactive MAPK14. Additinally, the results suggest that Pf is not a type I½or type II inhibitor of inactive MAPK14 and is unlikely to interfere with ATP to ADP conversion process needed for cell survival.

Further, we analyzed number of potential hydrogen bonds (Hbonds) fluctuations between Pf and the active and inactive MAPK14 proteins for 200 ns of the long 1 µs MD simulation (**Figure 4C**,**D)**. These results are consistent with the conclusions derived from the distance plots analysis. Specifically, for Pf simulations with the active conformation, we found that Pf mostly form 2 Hbonds during the course of the 200 ns of simulation time. In contrast, the results with inactive MAPK14 suggests that almost half the time Pf did not even indicate any Hbond formations suggesting poor Pf interactions with inactive MAPK14.

In summary, these MD simulation results suggest that Pf is stable at the catalytic binding site of active MAPK14 rather than that of inactive MAPK14 and has a higher preference towards the active conformation of MAPK14. Therefore, results indicate that Pf can potentially act as a Type 1 inhibitor of MAPK14. Type I inhibitors are defined as inhibitors bind to the active kinase conformation (DFG-Asp-in, αC-helix-in) and affect downstream signaling^26^.

### Potential Biological Pathway of Pf towards Improved Wound Healing

We have analyzed all our findings throughout the study and developed potential biological pathways shown in **Figure 5** which summarizes the related mechanistic pathways of Pf towards wound healing. The interactions of Pf were analyzed with several targets in the human proteome using a variety of methods: Pink colored proteins (MAPK14, MSK1, Akt1, FAK) were analyzed using molecular modeling techniques such as molecular docking and large-scale molecular dynamics (MD) simulations; purple colored protein (MAP3K4, MAP2K3, MAP2K6, MSK2, MAP2K2, ERK1, ERK2, PDK1) were predicted from the machine learning model; and grey colored proteins are not included in either analysis but shown to complete the pathway. Our study provides molecular and mechanistic basis to Pf’s targets that are involved in proliferation, inflammation, and cytoskeletal rearrangement/motility pathways towards wound healing.

**Figure 5.**
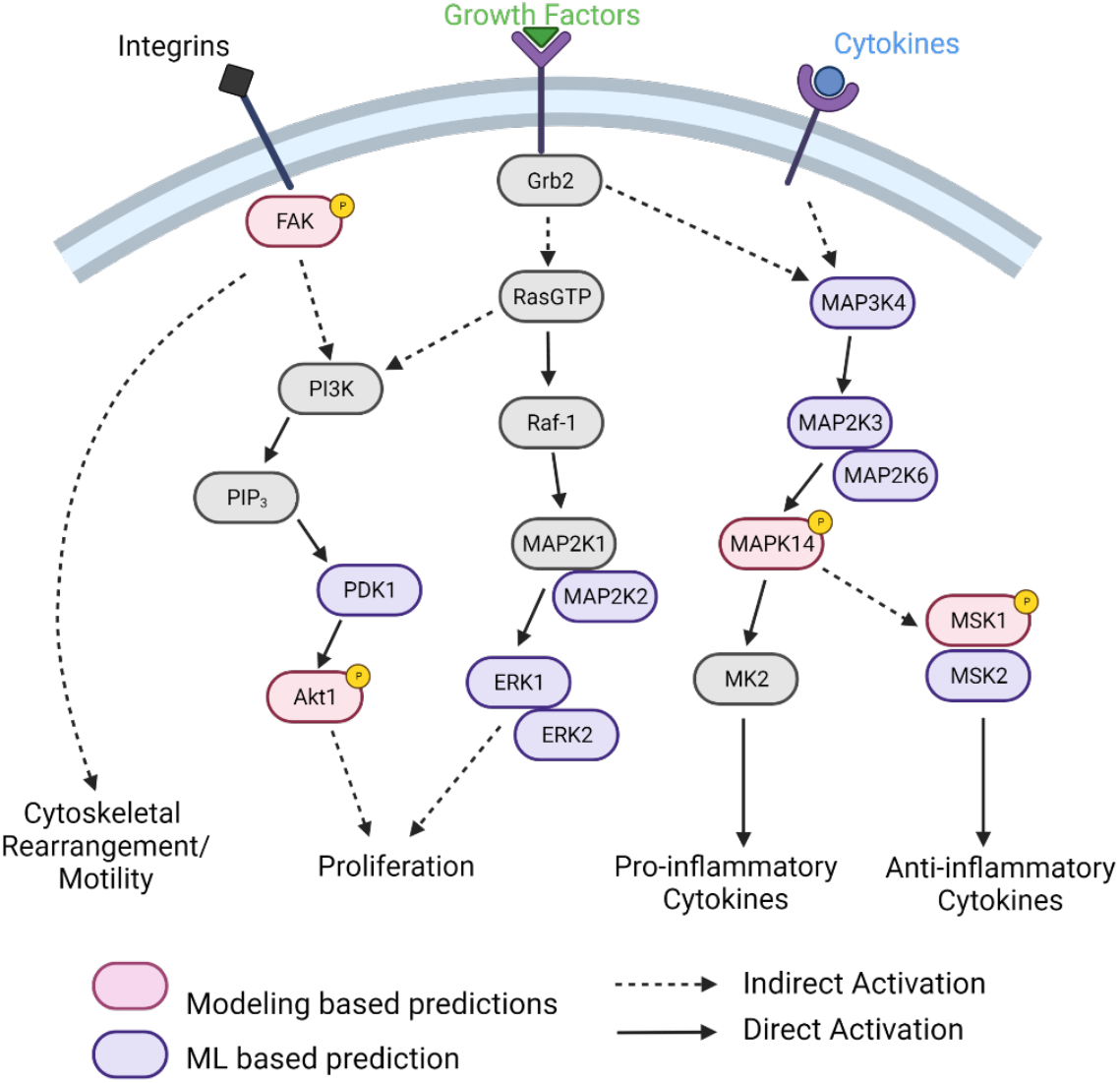
Biological pathways leading to regulate scar formation and potential biological targets of Pf identified using molecular modeling and machine learning.

## Conclusion

We have investigated Pf’s molecular mechanism using machine learning (**Figure 2**) and molecular modeling methods (**Figure 3-4**) and suggest a biological pathway model (**Figure 5**) to understand how it regulates phenomena such inflammation which lead to improved wound healing with reduced scarring. In addition to p38 MAPK, Akt1, and FAK as Pf targets that were derived from our prior *in vitro* and *in vivo* studies, using machine learning we discovered many other potential targets including MAP3K4, MAP2K3, MAP2K6, MSK2, MAP2K2, ERK1, ERK2, and PDK1 as part of the kinase signaling pathways that may contribute to the phenotype of Pf’s wound healing. In conclusion, we show that the ability of Pf in optimizing wound healing is not limited to targeting a single kinase or protein but could interact with multiple proteins with selected polypharmacological profile that is involved in kinase signaling, proliferation and inflammation resulting in improved wound healing with reduced scarring.

## Acknowledgements

This work was supported by the Geneva Foundation subaward no. S-1424-02 to G.C. originated from United States Department of Defense USAMRAA award W81XWH015-2-0083 to K.P.L. Additional funding, in part, by National Center for Advancing Translational Sciences ASPIRE Challenge and Reduction-to-Practice awards to G.C and the Purdue University Center for Cancer Research funded by NIH grant P30 CA023168 are also acknowledged. The content is solely the responsibility of the authors and does not necessarily represent the official views of the National Institutes of Health.

## Disclaimer

K.P.L. is a US Government employee. The views expressed are those of his and do not reflect the official policy or position of the U.S. Army Medical Department, Department of the Army, Department of Defense, or the U.S. Government.

