## Supporting Information for "Modeling Molecular Mechanisms of Pirfenidone Interaction with Kinases"

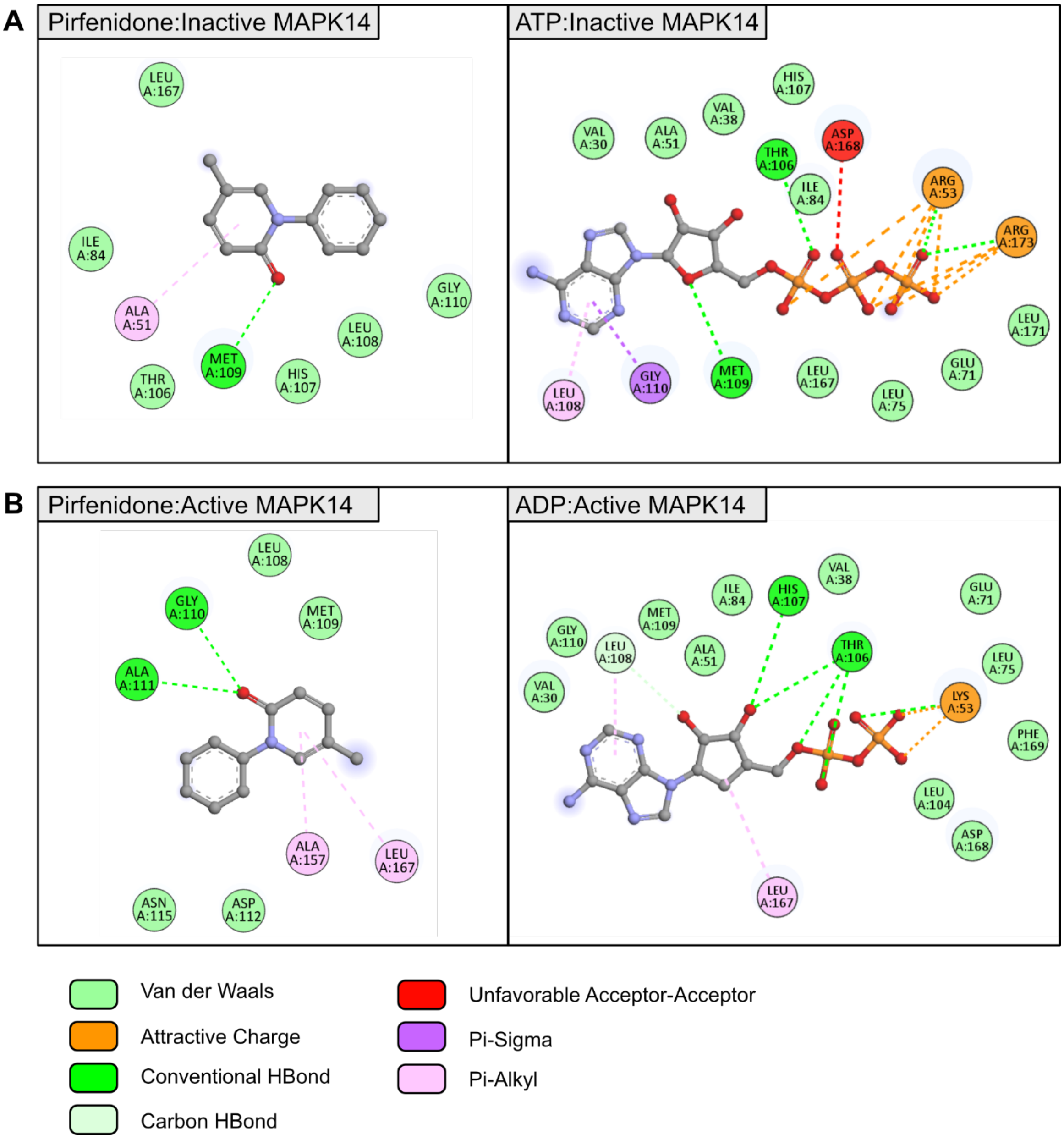


**Figure S1.** Interaction maps of Pf and ATP/ADP at their inactive/active MAPK14 binding sites. (A) 2D interaction maps of Pf and ATP with inactive MAPK14. (B) 2D interaction maps of Pf and ADP with active MAPK14.

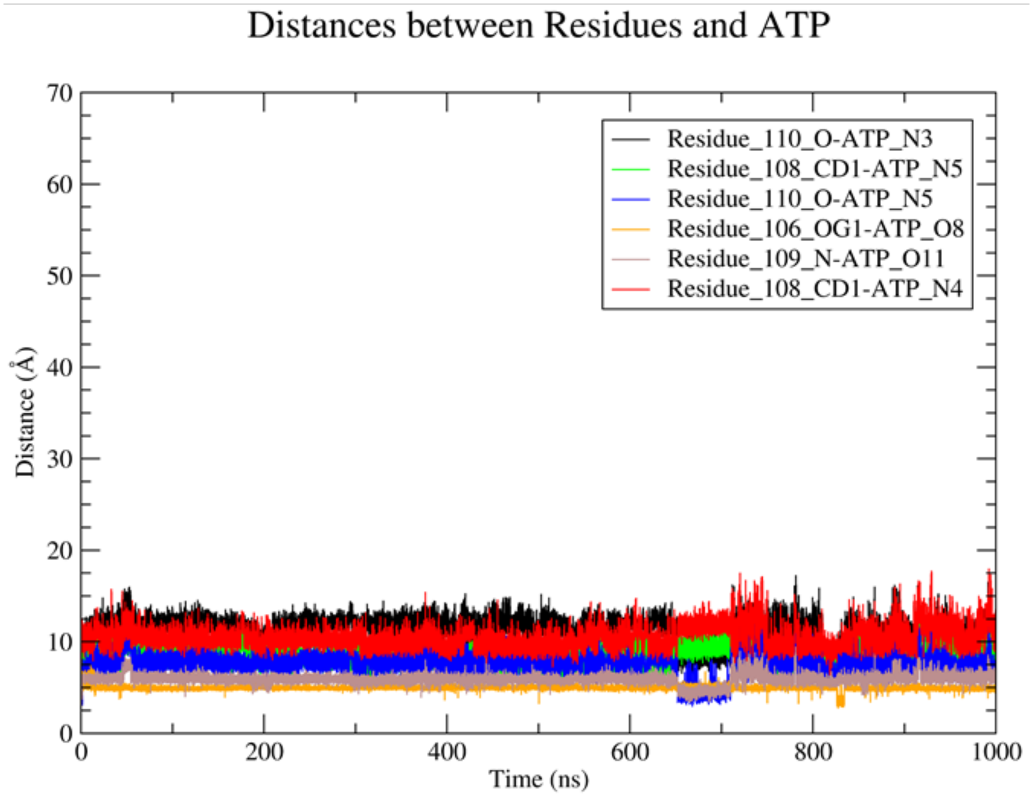


**Figure S2.** Distance trajectories between different atoms on the ATP and different hinge region residues on the MAPK14 for the inactive MAPK14 conformation.

**Table S1.** PDB IDs of all used p38-MAPK crystal structures. Proteins of *Mus musculus* are colored in blue and proteins of *Homo sapiens* are colored in red.

| P38-MAPK Protein | Active Conformations (PDB IDs) | Inactive Conformations (PDB IDs) |
|---|---|---|
| MAPK11 | - | 3gc8, 3gp0 |
| MAPK12 | 1cm8 | 6una |
| MAPK13 | 4myg | 3coi, 4eyj, 4eym, 4yno, 5ekn, 5eko |
| MAPK14 | 3py3 | 5eta, 5etc, 5etf, 5eti, 6sfo, 6y6v, 5nzz, 6y7w, 6y7x, 6y7y, 6y7z, 6y80, 6y81, 6y82, 6y85, 6y8h, 6ycu, 6ycw, 6yjc |

**Table S2.** Top five targets predicted for the pirfenidone molecule by the CPI machine learning model.

| **Name** | **Manning Name** | **HGNC Name** | **Kinase Name** | **Group** | **Family** | **Uniprot ID** | **ATP Binding** | **Pirfenidone Binding** | **PRF/ATP Ratio** |
|---|---|---|---|---|---|---|---|---|---|
| MAP2K3 | MAP2K3 | MAP2K3 | Dual specificity mitogen-activated protein kinase kinase 3 | STE | STE7 | P46734 | 0.92348 | 0.985849 | 1.06753 |
| MAP2K6 | MAP2K6 | MAP2K6 | Dual specificity mitogen-activated protein kinase kinase 6 | STE | STE7 | P52564 | 0.96943 | 0.993913 | 1.025247 |
| MSK2 | MSK2 | RPS6KA4 | Ribosomal protein S6 kinase alpha-4 | AGC | RSK | O75676 | 0.97614 | 0.998695 | 1.023106 |
| MAP2K2 | MAP2K2 | MAP2K2 | Dual specificity mitogen-activated protein kinase kinase 2 | STE | STE7 | P36507 | 0.29582 | 0.872099 | 2.947994 |
| Erk1 | Erk1 | MAPK3 | Mitogen-activated protein kinase 3 | CMGC | MAPK | P27361 | 0.44676 | 0.867337 | 1.941377 |
| Erk2 | Erk2 | MAPK1 | Mitogen-activated protein kinase 1 | CMGC | MAPK | P28482 | 0.41336 | 0.852424 | 2.062143 |

| | | | | | | | | | |
|---|---|---|---|---|---|---|---|---|---|
| PDK1 | PDK1 | PDPK1 | 3-phosphoinositide-dependent protein kinase 1 | AGC | PDK1 | O15530 | 0.25924 | 0.932434 | 3.596785 |
